# Motivations, concerns and selection biases when posting preprints: a survey of bioRxiv authors

**DOI:** 10.1101/2021.09.07.459259

**Authors:** Nicholas Fraser, Philipp Mayr, Isabella Peters

**Author notes:** Correspondence /.

## Abstract

Since 2013, the usage of preprints as a means of sharing research in biology has rapidly grown, in particular via the preprint server bioRxiv. Recent studies have found that journal articles that were previously posted to bioRxiv received a higher number of citations or mentions/shares on other online platforms compared to articles in the same journals that were not posted. However, the exact causal mechanism for this effect has not been established, and may in part be related to authors’ biases in the selection of articles that are chosen to be posted as preprints. We aimed to investigate this mechanism by conducting a mixed-methods survey of 1,444 authors of bioRxiv preprints, to investigate the reasons that they post or do not post certain articles as preprints, and to make comparisons between articles they choose to post and not post as preprints. We find that authors are most strongly motivated to post preprints to increase awareness of their work and increase the speed of its dissemination; conversely, the strongest reasons for not posting preprints centre around a lack of awareness of preprints and reluctance to publicly post work that has not undergone a peer review process. We additionally find weak evidence that authors preferentially select their highest quality, most novel or most significant research to post as preprints, however, authors retain an expectation that articles they post as preprints will receive more citations or be shared more widely online than articles not posted.

## Introduction

Preprints have become an integral part of the scholarly communication process. Whilst no singular authoritative definition of a “preprint” (in some disciplines referred to as a “working paper”) exists (Chiarelli et al., 2019; Tennant et al., 2018), they are commonly considered to be complete versions of scientific manuscripts that are posted to open online repositories prior to undergoing formal peer review and publication in a scientific journal. Posting of preprints therefore allows subversion of the traditional scientific publishing process; results are made available for dissemination immediately at the expense of journal- or conference-organised peer review and production, a process that can take between 9 and 18 months on average, dependent on the discipline (Björk & Solomon, 2013). In some scientific disciplines preprints have already become established as a norm. For example, in Physics and Mathematics, the preprint server arXiv (https://arxiv.org) has become widely established since its launch in 1991 (Ginsparg, 2011), with ~1.9 million preprints currently online (https://arxiv.org/stats/monthly_submissions), representing ~20% of the entire Physics and Mathematics journal literature available in the Web of Science (Larivière et al., 2014).

Experiments with preprints in the biological sciences date back to at least the early 1960s, when the National Institutes of Health (NIH) in the United States began circulating unreviewed preprints amongst Information Exchange Groups (IEGs) via physical postal services (Cobb, 2017). However, the scheme was abandoned in 1966 following pushback from publishers and journal editors who refused to publish articles that had been previously circulated via IEGs. Since then, a number of other attempts to bring preprints to the biological sciences were made, including e-BioMed (proposed in 1999, but never launched) (Varmus, 1999), ClinMed Netprints (active from 1999 to 2005) (Delamothe et al., 1999), and Nature Precedings (active from 2007 to 2012) (Nature, 2007). Yet, none of these endeavours stood the test of time, either as a result of continued lobbying by journal publishers, or weak uptake by authors; Nature Precedings only published 3,447 preprints in the 6 years it was active, and the announcement of closure noted that “the *Nature Precedings* site is unsustainable as it was originally conceived” (Nature, 2012). A more enduring effort to bring preprints to the biological sciences is through the quantitative biology (q-bio) section of arXiv, launched in 2003 and still active today. Even so, the uptake of q-bio has been relatively limited; only 17,322 preprints were published in q-bio between 2003 and 2019, representing <0.2% of the total biomedical literature available on PubMed (https://pubmed.ncbi.nlm.nih.gov/) during the same time interval.

In 2013, bioRxiv (https://biorxiv.org) was launched as a new preprint server for the biological sciences, hosted and operated by the Cold Spring Harbor Laboratory (CSHL) (Sever et al., 2019). In comparison to previous efforts to establish a dedicated biological preprint server, bioRxiv has been more successful: at the time of writing >130,000 preprints have been posted to bioRxiv, and submission rates continue to increase. The success of bioRxiv led CSHL to launch medRxiv (https://medrxiv.org), a “sister” preprint server aimed at the medical and health sciences, whilst a growing wave of other new preprint servers operated by non-profit academic groups and for-profit publishers now also cover aspects of biological and medical sciences (Kirkham et al., 2020). The rapid increase in the number and breadth of preprint servers in the past decade can be attributed to a number of factors, perhaps most importantly a swell in support from institutions and funders (e.g. the NIH “encourages investigators to use interim research products, such as preprints, to speed the dissemination and enhance the rigor of their work”; NIH, 2017) as well as increasing acceptance of preprints by journals, many of which now have partnerships established with preprint servers to support direct transfers of preprints to their journals (Penfold & Polka, 2020). In the last year, preprint servers including bioRxiv and medRxiv have experienced a surge of preprints in response to the COVID-19 pandemic; in the first 10 months of the pandemic, ~25% of all COVID-19 related biomedical literature was posted as preprints (Fraser et al., 2021). In comparison, only ~5% of literature related to Zika virus (2015-2017) and Western Africa Ebola virus (2014-2016) was posted as preprints, highlighting the recent shift in publication patterns towards preprint usage (Johansson et al., 2018).

Three recent studies have concluded that there exists a sizeable “advantage” for journal articles that were previously posted as preprints on bioRxiv, in terms of citations and various altmetric indicators (i.e. indicators of usage/sharing on online platforms) (Serghiou & Ioannidis, 2018; Fu & Hughey, 2019; Fraser et al., 2020). Two of these studies have additionally attempted to control for a number of external factors that may influence citation and altmetric counts, such as the number of authors, journal authority, and author demographics (e.g. author gender, country). Even when controlling for these factors, the size of the citation/altmetric advantage remains substantial: Fu and Hughey (2019) report that articles with a preprint received 1.36 times more citations and 1.49 times higher Altmetric Attention Scores than articles without a preprint, whilst Fraser et al. (2020) report that articles with a preprint received 1.56 times more citations, 2.33 times more tweets, 1.55 times more blog mentions, 1.47 times more mainstream media mentions, 1.30 times more Wikipedia citations and 1.81 times more Mendeley reads than articles without a preprint. These findings of a “bioRxiv citation/altmetric advantage” are in agreement with findings based on similar studies conducted on arXiv (Davis & Fromerth, 2007; Larivière et al., 2014; Moed, 2007; Wang, Chen, et al., 2020; Wang, Glänzel, et al., 2020), and related studies that have investigated the more general Open Access (OA) citation advantage, finding that OA articles tend to be more strongly cited than non-OA articles (Gargouri et al., 2010; Archambault et al., 2016; Piwowar et al., 2018).

Despite the large effect sizes of the bioRxiv citation/altmetric advantage reported, the aforementioned studies can still only be considered as observational in nature, i.e. it is not possible to ascribe the increase in citations or altmetrics to a causal mechanism. One key reason is that the studies only account for intrinsic authorship, article or journal properties, but not for factors relating to authors’ behaviour and decision-making processes that may lead to bias in the quality, and thus the “citeability” or “shareability” of preprints. This bias, previously termed the “Self-selection Bias Postulate”, or alternatively the “Quality Postulate” (Kurtz et al., 2005; Henneken et al., 2006; Moed, 2007; Davis & Fromerth, 2007; Gargouri et al., 2010), may itself be manifested in two dimensions: (1) authors may select their highest “quality” articles (where quality, in this sense, may refer to the articles that the authors believe will generate the highest citation/online impact) to post as preprints, which would consequently be more strongly cited or shared on online platforms than their lower quality articles, or (2) higher “quality” authors may preferentially post their articles as preprints compared to lower quality authors.

In this study, we aimed to primarily investigate the mechanism underlying the first dimension of the Self-selection Bias Postulate as applied to preprints, to investigate how author’s behaviours and motivations lead them to select certain articles to post as preprints. To achieve this, we conducted an online survey of bioRxiv preprint authors in which we asked them to report on their publishing and preprinting activities, with a specific focus on factors that support their decision to post or not post certain articles as preprints, and to report on differences between articles that they decide to post as preprints and those they do not (e.g. in terms of quality, novelty, or significance).

### Related work

A number of previous surveys have investigated the motivations and concerns of authors when posting or not posting preprints. A survey conducted by the Association for Computational Linguistics (ACL) (Foster et al., 2017) (N = 623) found that the two strongest motivations to post preprints, supported by 80% and 70% of the respondents who had posted a preprint, were to “publicize my research as soon as I think it is ready” and to “timestamp the ideas in the paper”, respectively. Only 32% of respondents reported that they have posted a preprint to “maximize the paper’s citation count”, implying that increasing a paper’s impact ranks relatively low amongst computational linguists’ motivations. Conversely, when respondents who had not posted a preprint were asked for their reasons for not doing so, 71% reported that there “was no need when I intend to publish my papers”, and 59% reported that they did not do so “to preserve the integrity of double-blind reviewing”.

A survey of community members of the Special Interest Group on Information Retrieval (SIGIR) (Kelly, 2018) (N = 159) found broadly similar results to that of the ACL survey: the two strongest motivations for posting preprints remained to “publicize my research as soon as I think it is ready” (67%) and to “timestamp the ideas in the paper” (81%), with only 48% of respondents posting preprints to “maximize the paper’s citation count”, whilst the most-agreed with reasons for not posting preprints were “I want to preserve the integrity of double-blind reviewing” (65%), and “I do not see the need when I intend to publish my papers at a conference or in a journal” (63%).

In the context of bioRxiv, Sever et al. (2019) previously conducted a survey of bioRxiv authors, readers, and non-users to ask about their motivations for posting preprints, reasons that they had not posted a preprint on bioRxiv (if applicable), and how posting a preprint on bioRxiv has benefited their careers. They found that the strongest motivations of preprint authors (N = 3,364) were to “increase awareness of your research” (80% of authors) and “to benefit science” (69%), whilst only 54% of authors did so “to stake a priority claim on your research”, a lower proportion than was reported in the respective questions of the ACL and SIGIR surveys. Unlike the ACL and SIGIR surveys, the survey of Sever et al. (2019) did not specifically ask authors about the effect of posting preprints on impact indicators, e.g. citations or altmetrics. Of those who had not posted a preprint to bioRxiv (N = 844), the most frequently reported reason was that authors “do not have enough data yet for a research manuscript” (35%), followed by “you are still deciding whether posting a preprint is the right choice for you” (24%), although only 10% of bioRxiv non-authors agreed that they “are not convinced that preprints are a good idea”. In terms of benefits of posting a preprint to bioRxiv, by far the most-frequently agreed with reason was that they “Increased awareness of your research” (73%) - the second most-frequently agreed with reason was that they “Helped stake a priority claim for your research” (28%).

A more recent survey conducted by ASAPbio (https://asapbio.org/), a nonprofit that advocates for “innovation and transparency in life sciences communication”, also investigated the perceived benefits and concerns surrounding preprints (N = 546) (ASAPbio, 2020). The preliminary findings broadly echoed those of Sever et al. (2019): the benefit most strongly agreed with was “increasing the speed of research communication”; however, agreement was stronger amongst authors who had previously posted preprints versus those who had not. Although all of these surveys have targeted different groups at different times, some clear themes have emerged in terms of factors that support authors’ decision to post preprints: they do so primarily to increase awareness of their work (i.e. increased distribution and/or readership amongst colleagues and other scientists), to increase the speed of its dissemination into relevant communities, to enable free/unrestricted access for readers, to receive more feedback, and to stake a priority claim on their research ideas, whilst other motivating factors such as increasing citation counts, or supporting career development (e.g. to cite a preprint in a job application) appear to be secondary. In contrast, factors that discourage authors’ from posting preprints focus mainly on concerns surrounding preprint integrity due to lack of peer-review, risks of premature reporting in the media, and of work being “scooped” when published too early. Notably, external pressures to post or not post preprints (e.g. posting due to institutional/funding policies, or not posting due to journal policies/the Ingelfinger rule^1^) appear to rank relatively low amongst researchers’ motivations or concerns when deciding whether to post a preprint.

A related study by Chiarelli et al. (2019) also explored the perceived benefits and challenges of preprint posting, although their study concentrated on a range of stakeholders not limited to preprint authors (e.g. research funders, preprint servers and service providers), and used semi-structured interview methods rather than quantitative surveys. However, the results largely echoed those of the previously discussed surveys: stakeholders reported the most important benefits of preprints to be early and rapid dissemination, and increased opportunities for feedback (both mentioned by >20 interviewees from a sample of 38 interviewees), whilst increasing citation counts was a less important factor (mentioned by <10 interviewees). The most important challenges of preprints were the lack of quality assurance, limited use of commenting/feedback, risk of media reporting incorrect research, possible harm in the case of sensitive areas, and questionable value of self-appointed reviewers (all mentioned by >20 interviewees). The study concludes that a one-size-fits-all approach to preprints is not feasible: different disciplinary communities are at different stages in their processes of using, adopting or experimenting with preprints, and that the future of preprint servers and related infrastructure remains unclear.

### Motivation and Research Gap

Our survey approach contains some similarities to previous surveys that have investigated the motivations and concerns of authors to post or not post preprints, but differs in several important aspects:

1. We ask authors about their motivations and concerns when posting or not posting preprints, with a specific focus on articles that were eventually published in scientific journals.
2. We ask authors to make direct comparisons between their published journal articles that were and were not posted as preprints.

Focusing on these missing aspects from previous surveys will allow us to partially fill the gap in understanding the Self-selection Bias Postulate: we hope to be able to understand the reasons that authors are motivated to post or not post certain journal articles as preprints, and relate these reasons to factors such as quality, novelty and expected citation performance of the articles.

Our previous work also showed that citation/online sharing rates of journal articles posted as preprints are related to various author-specific factors, e.g. higher citation rates were associated with more senior first and last authors, male first authors, and first authors from the USA (Fraser et al., 2020). We therefore aimed to distinguish differences in motivations and selection strategies of authors to post preprints between multiple demographics groups, namely the participants’ country of residence (US versus non-US), gender and career status.

## Methods

### Survey participants

Our pool of potential survey participants consisted of corresponding authors of preprints posted to bioRxiv between November 2013 (coinciding with the launch of bioRxiv) and December 2018. Email addresses of corresponding authors were harvested by crawling of bioRxiv public web pages using the R packages *rcrossref* for harvesting preprint DOIs (Chamberlain et al., 2020) and *rvest* for reading and extracting HTML (Wickham, 2020). For each preprint, corresponding author email addresses were extracted from the HTML meta tag “citation_author_email” (note that a single preprint can list multiple corresponding author email addresses). We harvested a total of 49,447 raw email addresses associated with 23,986 preprints. Email addresses were subsequently validated through regular expressions, and duplicate email addresses were removed – these were either the result of authors submitting multiple preprints, or in a small number of cases the same email address being listed multiple times on the same preprint. Following these steps, we produced a list of 24,633 unique validated email addresses. Potential participants were invited via email to participate in the survey on 27^th^ & 28^th^ February 2020, and a follow-up invitation was sent on 25^th^ March 2020. Responses were collected until 29^th^ April 2020. Of 1,847 participants who began the survey, 1,444 participants fully completed it, representing a drop-out rate of 27.9% and a response rate (for fully complete surveys) of 5.9%. In all the following analysis, we only consider data from participants who fully completed the survey.

### Survey design and development

A generalised overview of the survey design is shown in **Figure 1**. The survey was built with LimeSurvey version 2.73.1, and hosted on a server maintained by the University of Kiel, Germany.

**Figure 1:**
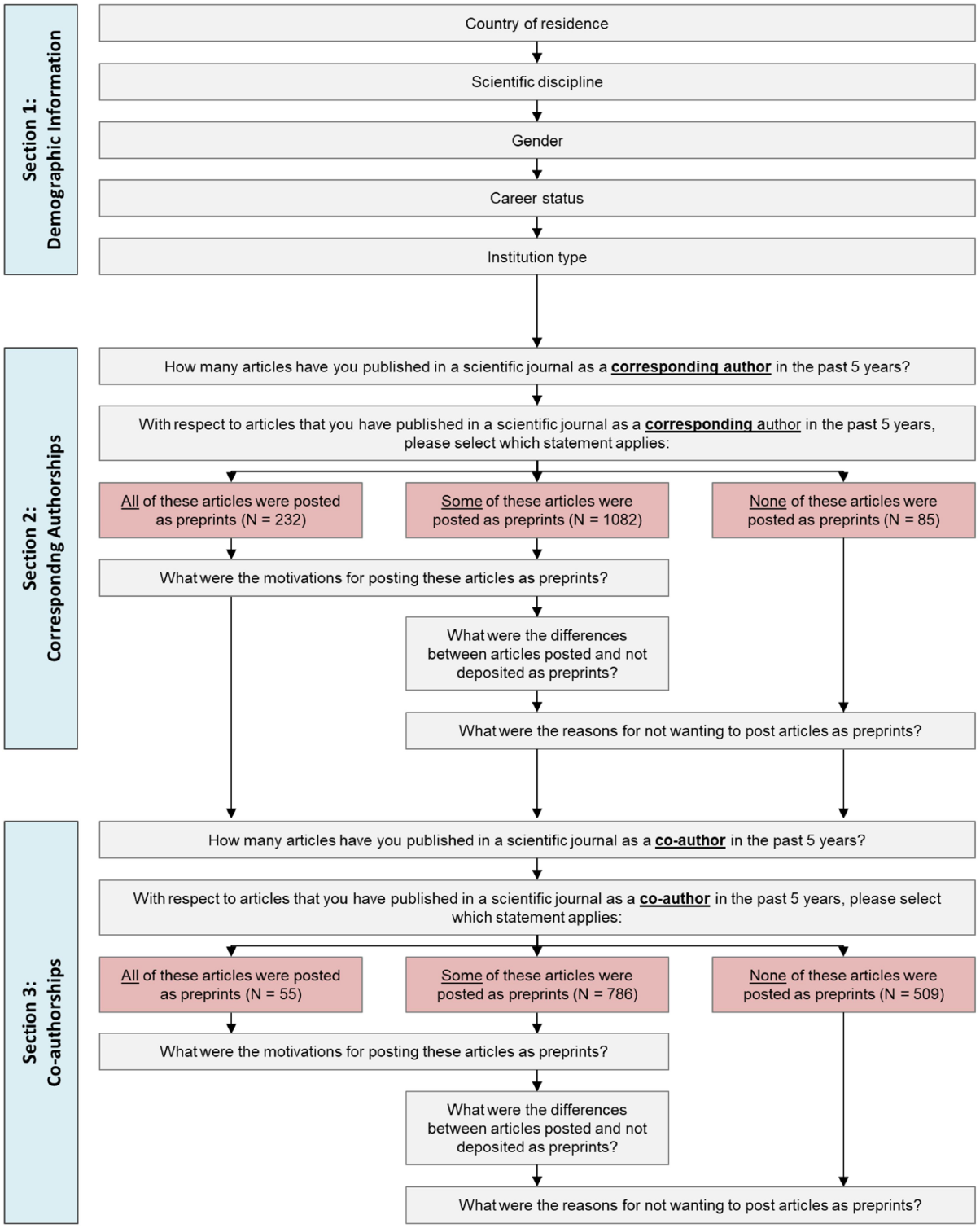
Generalized overview of survey design.

In brief terms, the survey was divided into three main sections:

The first section asked participants to report their demographic background (country of residence, main scientific disciplines, gender, career status and the institution type of their employer). For each question, participants were also able to opt-out of answering.

The second section asked participants to consider their recent (past 5 years) record of publishing in scientific journals as a *corresponding author*. With respect to journal articles published in that time period, participants were grouped according to the proportion of those articles that were also posted as preprints (authors who reported that they did not publish any journal articles in that time were excluded), and then directed conditionally to the following survey questions: authors who reported posting all of their journal articles as preprints were surveyed on their motivations for posting preprints (N = 232), authors who reported that none of their journal articles were posted as preprints were surveyed on their reasons for not posting preprints (N = 85), and authors who reported that a selection of their journal articles were posted as preprints were surveyed on both of the above areas, as well as self-reported differences between journal articles that were and were not posted as preprints (for example, whether there was a difference in the quality or novelty of articles that were and were not posted as preprints) (N = 1,082).

The third section of the survey repeated the second section in structure, but asked respondents to instead focus on their record of publishing in scientific journals as a *co-author*, defined in the survey as articles where the author was in any position other than the corresponding author (N = 55, 786 and 509 for authors reporting that all, some or none of their co-authored articles were posted as preprints, respectively). This division between sections 2 and 3 was designed to determine if differences arose for articles where authors had a higher degree of autonomy over the publication strategy (i.e. as a corresponding author), versus those where authors’ responsibility for the publication strategy is perhaps more passive or moderated by the decisions of the leading authors.

### Regression Analysis

To better understand potential differences in preprinting behaviour between demographic groups (namely gender, career status and country of residence), we conducted ordinal logistic regression for each of our survey questions that were answered on a Likert scale. To account for the high granularity of participant information, for the independent variable of “career status” participants were pooled into “Early Career” (Master’s students, PhD students, Postdoctoral Researchers) or “Late Career” (Assistant/Associate/Full Professor) groups, and for the independent variable “country of residence” into “US” or “non-US” groups (this pooling is derived from previous work that has shown that ~50% of first and last authors of bioRxiv preprints are from the US (Fraser et al., 2020)). Participants that chose not to report certain demographic information were excluded from the regression analysis, as were participants with the “career status” of business or healthcare professionals, due to low numbers of participants from these groups. Regression analysis was limited to responses based on participants’ experiences as a corresponding author (i.e. Section 2 from **Figure 1**). Regression analysis was conducted using the *polr()* function from the R package *MASS* (Venables & Ripley, 2002).

All regression results are presented as exponentiated odds-ratios (OR), with 95% confidence intervals. Odds-ratios of an ordinal logistic regression model can generally be interpreted as the odds that a one-unit increase in an independent variable is associated with a higher value on an ordinal dependent variable: an OR greater than 1 indicates a positive association and an OR less than 1 indicates a negative association. In practical terms for our study (where we only use binary categorical independent variables), odds-ratios are therefore interpreted as the odds that the reported group (e.g. “early career” in the independent variable “career status”) will rank an answer one-point higher on a Likert scale (e.g. “strongly agree” instead of “agree”) than the comparison group (“late career”).

### Analysis of free-text comments

For survey questions relating to motivations, concerns and differences in preprint posting behaviour in sections 2 and 3, participants were provided with the opportunity to add free-text responses to expand on their answers or add any further context. Free-text comments were coded following the general inductive approach of Thomas (2006): comments were initially read to gain familiarity with their contents, and then given one or several unstructured labels reflecting their contents (e.g. the comment *“Preprints allowed everyone to read the manuscript when I could not publish them Open Access, due to prohibitive costs of open access fees.”* was initially given the labels “accessibility” and “cost”). Labels were subsequently iteratively revised, grouped and refined into ~8-10 broader categories (e.g. “accessibility” and “cost” labels were subsequently grouped under the category “open science”) that conveyed the core themes of the comments. A small number of responses that did not contain information relevant to the survey question asked were ignored. Labels that did not fit within any of the broader categories and occurred <10 times across all free-text responses were grouped into the category “other”. All coding was conducted by the first author of this study, NF. A codebook was maintained during the coding process, with names, descriptions and examples of each theme. A total of 1,095 comments were coded following this process: 291 related to motivations for posting preprints, 317 related to reasons for not posting preprints, and 487 to differences between articles that were and were not posted as preprints.

### Data Storage and Processing

Data collected in this study, as well as the survey design and LimeSurvey files, are available on GitHub (https://github.com/nicholasmfraser/biorxiv_survey) and archived on Zenodo (https://zenodo.org/10.5281/zenodo.5166749). Raw free-text responses and participant email addresses were removed from the final archived datasets due to the potential for participant identification. Following collection of the survey data and coding of free-text responses, all subsequent analysis and visualizations were produced with R version 4.0.1 (R Core Team, 2020).

## Results

### Survey participants

An overview of demographic information for survey participants is shown in **Figure 2**. In general terms, survey participants were biased towards those based in North America and Europe, to male participants, and to researchers based at universities.

**Figure 2:**
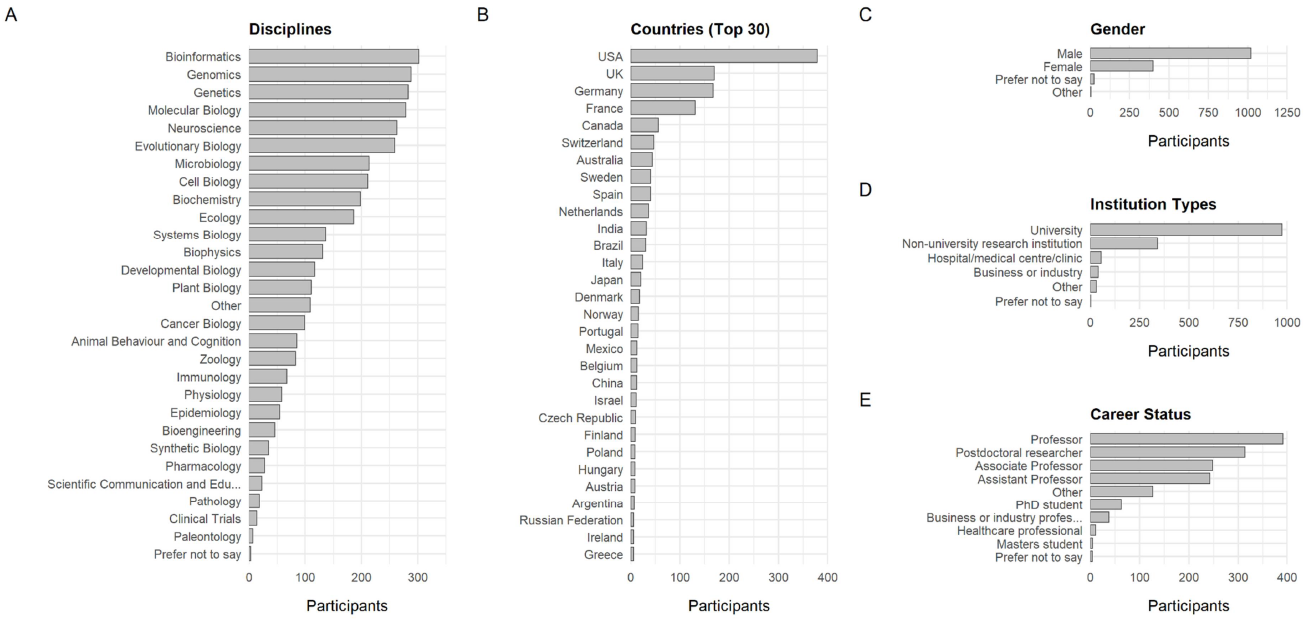
Demographics of survey participants (N=1,444). (A) Main scientific disciplines. Note that disciplines correspond to bioRxiv submission categories, and participants could select multiple disciplines. (B) Country of residence. The top 30 countries by total participants are shown. (C) Gender. (D) Institution type. (E) Career status.

Observed biases in the demographics of survey participants is at least partially a result of the bias in demographics of authors posting preprints to bioRxiv who were contacted to take the survey. For example, the most strongly represented disciplines of Bioinformatics, Genomics and Genetics amongst survey participants are also amongst the disciplines with the highest number of preprints (Abdill & Blekhman, 2019), whilst previous studies have also found that authors from the US are overrepresented on bioRxiv (Fraser et al., 2020; Abdill et al., 2020), as are male authors (Fraser et al., 2020). However, there also exist some discrepancies in participant representation compared to our expectations; for example, only 12 of 1444 participants (0.8%) are from China, whilst 3.6% of last authors and 6.5% of all authors of bioRxiv preprints are from China (Abdill et al., 2020).

### Publishing Behaviour

Participants were asked to report on their publishing behaviour over the past 5 years, including the number of articles they have published in peer-reviewed scientific journals in that time period (both as a corresponding author and as a co-author), and the proportion of those articles that were posted as preprints (simplified to the categories of “All”, “Some” or “None”) (**Figure 3**).

**Figure 3:**
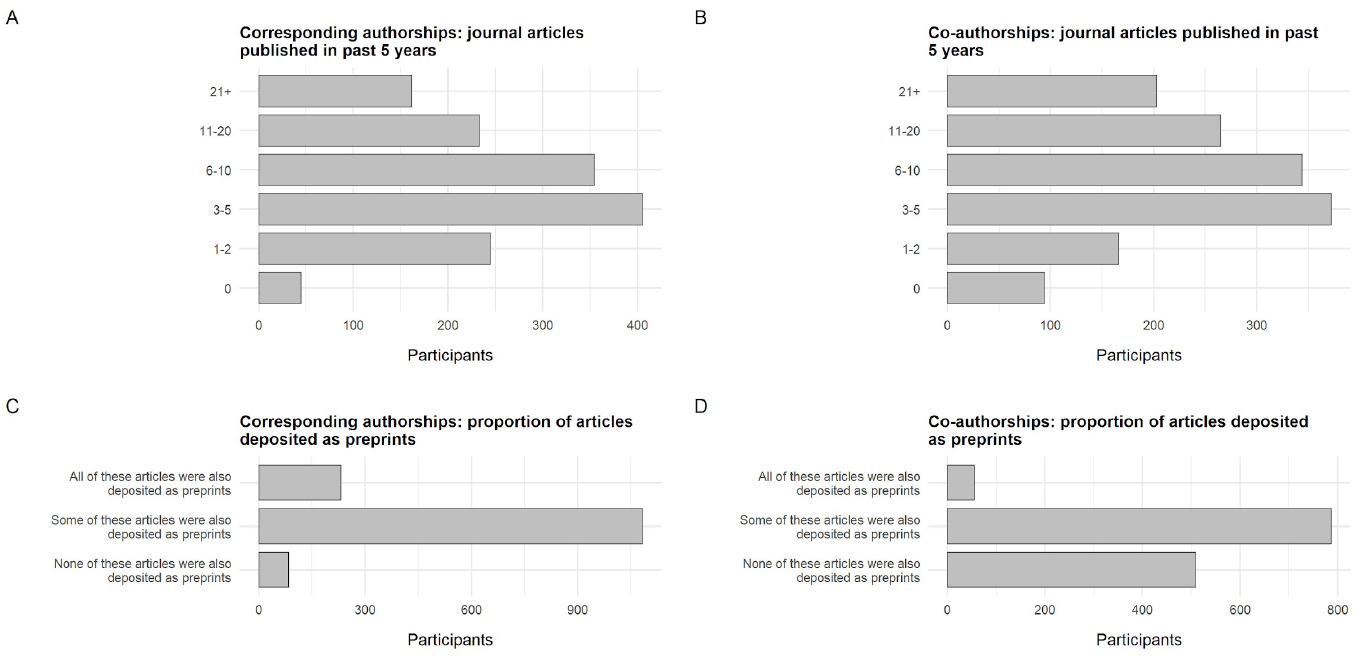
Publishing behaviour of survey participants over the past 5 years. (A) Number of articles published by survey participants in peer-reviewed scientific journals in the past 5 years *as a corresponding author*. (B) Number of articles published by survey participants in peer-reviewed scientific journals in the past 5 years *as a co-author*. (C) Proportion of articles in (A) that were posted as preprints. (D) Proportion of articles in (B) that were posted as preprints. Authors who reported that they published 0 articles in the past 5 years as a corresponding author or as a co-author were not included in (C) or (D), respectively.

Similar patterns of publication behaviour were found for corresponding authorships and coauthorships – the most frequent category for both groups was 3-5 publications (28.0% for corresponding authorships vs 25.8% for co-authorships). At the extremes, a higher proportion of participants reported that they had published zero articles as a co-author compared to as a corresponding author (6.5% vs 3.1%), and a higher proportion reported that they had published 21 + articles as a co-author compared to as a corresponding author (14.0% vs 11.2%). In terms of preprinting behaviour, the majority of participants reported that they only posted some of their articles as preprints during that time period both as a corresponding author (77.3%) and as a co-author (58.2%), whilst a higher proportion of participants reported posting all of their articles as a corresponding author as preprints (16.6%) compared to their articles as a co-author (4.1%).

### What motivates researchers to post preprints?

Survey participants who reported that they had posted all or some of their recent journal articles as preprints (**Figure 3**) were presented with questions that focused specifically on this set of articles. Questions covered three main focus areas: decision-making (who was responsible for the decision to post a preprint) (**Figure 4; Table 1**), motivating factors (what internal/external factors made the authors want to post a preprint) (**Figure 5; Table 2**), and the benefits received in terms of article citation/online impact (**Figure 6, Table 3**). Participants were also provided with a free-text area to expand on their answers for this section or add further reasoning (**Table 4**).

**Figure 4:**
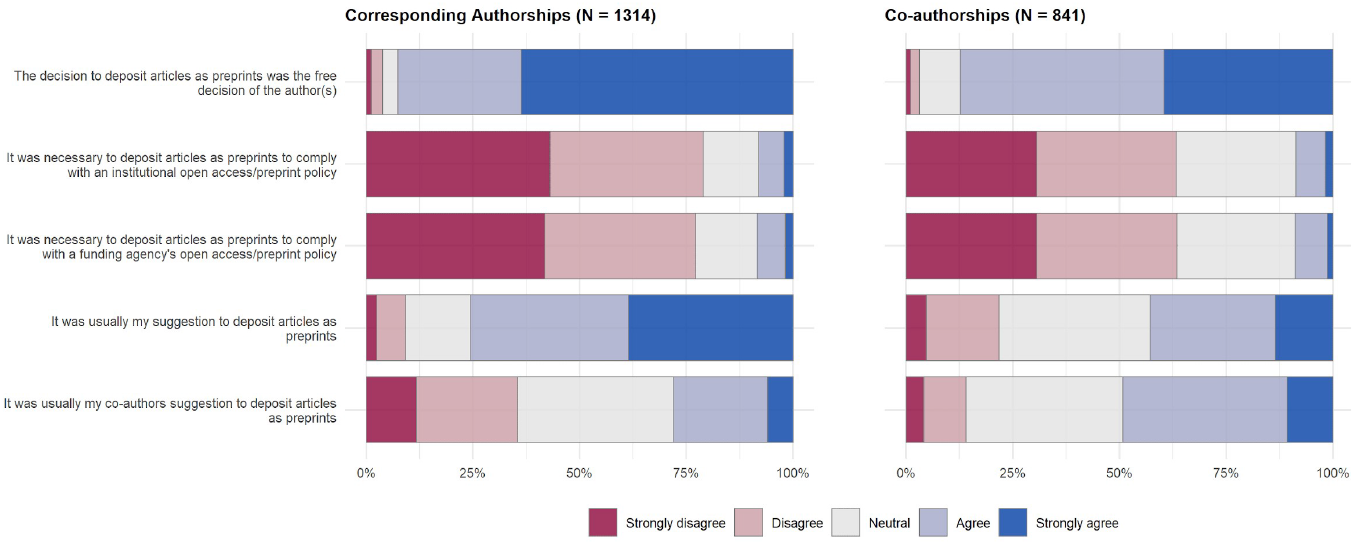
Decision-making for posted preprints. Results show answers to survey questions on a 5-point Likert scale, divided by articles that authors published as a corresponding author (left) and articles that authors published as a co-author (right).

**Table 1:** Results of ordinal logistic regression for survey questions in Figure 4 (corresponding authorships only; N = 1146). Values are presented as exponentiated odds-ratios, with the range in parentheses representing 95% confidence intervals. Significant results at the 95% level are indicated in **bold** and with “*”.

| Survey Question | Female | Early Career | non-US |
| --- | --- | --- | --- |
| The decision to post articles as preprints was the free decision of the author(s) | 0.979<br>(0.751 - 1.280) | 1.080<br>(0.830 - 1.410) | **0.757 \***<br>**(0.570 - 0.999)** |
| It was necessary to post articles as preprints to comply with an institutional open access/preprint policy | 1.249<br>(0.981 - 1.591) | **0.683 \***<br>**(0.534 - 0.872)** | **1.407 \***<br>**(1.094 - 1.813)** |
| It was necessary to post articles as preprints to comply with a funding agency's open access/preprint policy | 1.279<br>(1.007 - 1.626) | **0.692 \***<br>**(0.542 - 0.881)** | 1.225<br>(0.956 - 1.574) |
| It was usually my suggestion to post articles as preprints | **0.680 \***<br>**(0.533 - 0.867)** | 0.948<br>(0.748 - 1.204) | 0.991<br>(0.772 - 1.271) |
| It was usually my co-authors suggestion to post articles as preprints | 1.266<br>(0.998 - 1.607) | **1.317 \***<br>**(1.039 - 1.668)** | 0.842<br>(0.661 - 1.071) |

In terms of decision-making (**Figure 4**), participants overwhelmingly reported that the decision to post their articles as preprints was the free decision of the authors (92.6% agreed/strongly agreed for corresponding authorships, 87.4% for co-authorships), and was usually not externally driven by open access/preprint policies of their institutions (78.9% disagreed/strongly disagreed for corresponding authorships, 63.4% for co-authorships) or funding agencies (77.2% disagreed/strongly disagreed for corresponding authorships, 63.5% for co-authorships). For articles where participants acted as the corresponding author, a higher proportion reported that they themselves suggested to post an article as a preprint (75.6% agreed/strongly disagreed) than their co-authors (28.0% agreed/strongly agreed); the effect is reversed for articles where participants acted as a co-author, where a higher proportion reported that their co-authors made the suggestion (49.2%) compared to making the suggestion themselves (42.9%).

Regression analysis for the set of questions presented in **Figure 4 (Table 1)** show that non-US authors were less likely to have made the decision to post preprints freely (OR: 0.757, 95% CI: 0.570 - 0.999) compared to their US counterparts. Early career researchers were less likely to post preprints to comply with open access/preprint policies from their institution (OR: 0.683, 95% CI: 0.534 - 0.872) or funding agency (OR: 0.692, 95% CI: 0.542 - 0.881), whilst non-US authors were more likely to post preprints to comply with an institutional open access/preprint policies from their institution (OR: 1.407, 95% CI: 1.094 - 1.813). Female participants were less likely to have suggested posting articles as preprints (OR: 0.680, 95% CI: 0.533 - 0.867) than male participants, whilst early career participants were more likely to have had a co-author who suggested posting articles as preprints (OR: 1.317, 95% CI: 1.039 - 1.668).

Following questions surrounding decision making, survey participants were asked about their own motivations to post preprints; a selection of motivating factors were chosen based on commonly occurring themes from previous survey results (Foster et al., 2017; Kelly, 2018; Sever et al., 2019): to increase awareness of their work, to claim priority over results, to benefit the scientific enterprise, to increase the amount of feedback received, or to increase rate of dissemination (**Figure 5**).

**Figure 5:**
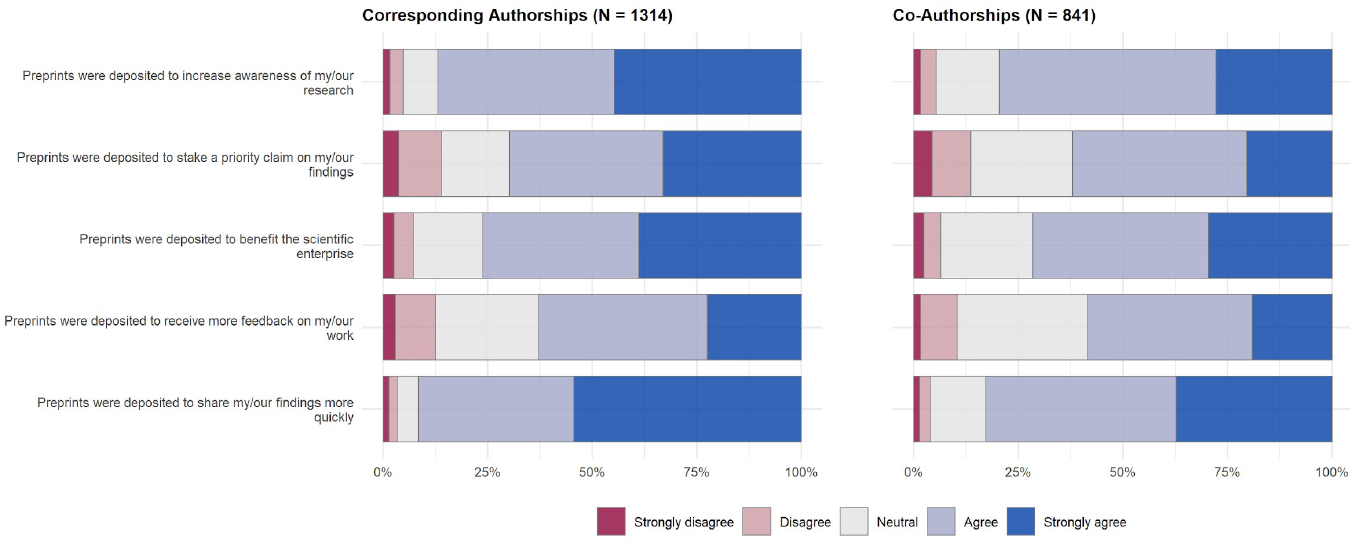
Motivations for posting preprints. Results show answers to survey questions on a 5-point Likert scale, divided by articles that authors published as a corresponding author (left) and articles that authors published as a co-author (right).

In general, participants reported that all of these factors positively motivated them to post preprints; the strongest motivator was to share their findings more quickly (91.5% agreed/strongly agreed for corresponding authorships, 82.7% for co-authorships), followed by increasing awareness of their research (86.9% agreed/strongly agreed for corresponding authorships, 79.5% for co-authorships).

Receiving more feedback on their work was the weakest motivator overall (62.8% agreed/strongly agreed for corresponding authorships, 58.5% for co-authorships), yet remained a motivating factor for the majority of authors. In general, motivating factors do not appear to differ strongly between articles that authors acted as a corresponding author versus those where they acted solely as a co-author.

Regression analysis for the set of questions presented in **Figure 5 (Table 2)** show that early career researchers are motivated more strongly to post preprints to increase aware of their research (OR: 1.317, 95% CI: 1.030 - 1.687) and to receive more feedback on their work (OR: 1.367, 95% CI: 1.079 - 1.732), but less likely to post preprints to stake a priority claim on their work (OR: 0.668, 95% CI: 0.528 - 0.846). Non-US authors were less likely to post preprints to increase awareness of their research (OR: 0.679, 95% CI: 0.524 - 0.878), to benefit the scientific enterprise (OR: 0.505, 95% CI: 0.392 - 0.651) or to share their findings more quickly (OR: 0.711, 95% CI: 0.543 - 0.927). These results are in agreement with results from the previous section, where non-US authors reported less freedom in decision-making over posting their preprints, and posted preprints more often to comply with institutional open access/funding policies; non-US authors may therefore be less motivated by external factors when deciding whether to post preprints or not.

**Table 2:** Results of ordinal logistic regression for survey questions in Figure 5 (corresponding authorships only; N = 1146). Values are presented as exponentiated odds-ratios, with the range in parentheses representing 95% confidence intervals. Significant results at the 95% level are indicated in **bold** and with “*”.

| Survey Question | Female | Early Career | non-US |
| --- | --- | --- | --- |
| Preprints were posted to increase awareness of my/our research | 1.041<br>(0.812 - 1.336) | **1.317 \***<br>**(1.030 - 1.687)** | **0.679 \***<br>**(0.524 - 0.878)** |
| Preprints were posted to stake a priority claim on my/our findings | 0.995<br>(0.785 - 1.261) | **0.668 \***<br>**(0.528 - 0.846)** | 0.924<br>(0.721 - 1.183) |
| Preprints were posted to benefit the scientific enterprise | 0.805<br>(0.631 - 1.027) | 0.873<br>(0.688 - 1.108) | **0.505 \***<br>**(0.392 - 0.651)** |
| Preprints were posted to receive more feedback on my/our work | 0.985<br>(0.777 - 1.249) | **1.367 \***<br>**(1.079 - 1.732)** | 0.956<br>(0.748 - 1.223) |
| Preprints were posted to share my/our findings more quickly | 1.183<br>(0.915 - 1.533) | 1.226<br>(0.951 - 1.583) | **0.711 \***<br>**(0.543 - 0.927)** |

The final questions in this section of the survey related to the potential benefits of preprints, in terms of increased citations and other forms of online dissemination (e.g. sharing on social media) (**Figure 6**). The largest group of participants responded neutrally with respect to the effect of posting preprints on citations (50.6% neutral for corresponding authorships, 48.9% for co-authorships), although a larger proportion of participants agreed/strongly agreed that posting preprints had positive benefits on citations (38.7% for corresponding authorships, 42.5% for co-authorships) than disagreed/strongly disagreed (10.6% for corresponding authorships, 8.6% for co-authorships). In contrast, the majority of participants reported positive effects of posting preprints on other forms of online dissemination (66.7% agreed/strongly agreed for corresponding authorships, 64.5% for co-authorships), suggesting that participants believe posting preprints has a stronger effect on other forms of online dissemination than on their citation impact.

**Figure 6:**
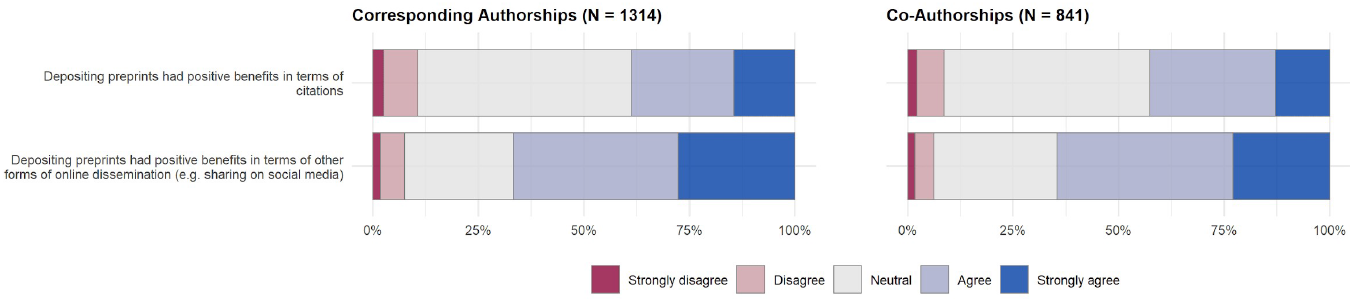
Benefits of posting preprints on article impact. Results show answers to survey questions on a 5-point Likert scale, divided by articles that authors published as a corresponding author (left) and articles that authors published as a co-author (right).

Regression analysis for the set of questions presented in **Figure 6 (Table 3)** show that early career researchers are more likely to report positive benefits of posting preprints both in terms of citations (OR: 1.318; 95% CI: 1.033 - 1.681) and other forms of online dissemination (OR: 1.318; 95% CI: 1.040 - 1.672), whilst non-US authors were less likely to report positive benefits in terms of other forms of online dissemination (OR: 0.637; 95% CI: 0.498 - 0.815).

**Table 3:** Results of ordinal logistic regression for survey questions in Figure 6 (corresponding authorships only; N = 1146). Values are presented as exponentiated odds-ratios, with the range in parentheses representing 95% confidence intervals. Significant results at the 95% level are indicated in **bold** and with “*”.

| Survey Question | Female | Early Career | non-US |
| --- | --- | --- | --- |
| posting preprints had positive benefits in terms of citations | 0.920<br>(0.720 - 1.176) | **1.318 \***<br>**(1.033 - 1.681)** | 0.797<br>(0.620 - 1.025) |
| posting preprints had positive benefits in terms of other forms of online dissemination (e.g. sharing on social media) | 1.077<br>(0.849 - 1.368) | **1.318 \***<br>**(1.040 - 1.672)** | **0.637 \***<br>**(0.498 - 0.815)** |

In addition to the above questions, respondents were provided with a free-text box in this section of the survey, in which they could write comments to elaborate on their motivations for posting their articles as preprints (**Table 4**). Free-text responses were coded into categories that represented the core themes of the free-text comments (plus a category “other” for comments that did not fit into one of the main categories). Note that one comment could belong to several categories.

**Table 4:** Classification of free-text responses, in response to the question “Were there any additional reasons that motivated you to deposit these articles as preprints?” Categories are ordered alphabetically (except for “Other”). Percentages for corresponding and co-authorships refer to the proportion of all responses containing the relevant category; responses could contain multiple categories (N = 253 comments on corresponding authorships, 38 comments on co-authorships).

| Category | Description | Examples | Corresponding authorship (%) | Co-authorship (%) |
| --- | --- | --- | --- | --- |
| Career development | To show evidence of research results in support of job/tenure/grant/graduate school applications; to add research outputs that do not intend to be published in a journal to a CV. | *“Yes, it gives additional weightage to your CV for people who don't have enough publications and looking for a new academic jobs.”*<br>*“The preprint was used as evidence of a high quality manuscript when applying for promotion”* | 36.8 | 29.7 |
| Competition | To claim priority on research findings; to prevent "scooping" of results by competitors; to allow publication of results concomitantly with presentations at conferences. | *“To coincide with conference presentations (hot off the presses results).”*<br>*“I had a bad experience with a paper that was scooped while our data had been submitted to several journals since two years.”* | 8.3 | 5.4 |
| Dissemination | To increase dissemination of work into relevant communities; to increase citation impact; to allow citation of work not published in journals by other researchers or by follow-up work. | *“Preprint can be cited in a second, follow-up article, before the first one is published. It is a better option than citing related research (conducted by our group) as “in press”.”* | 20.2 | 5.4 |
| Editorial process | Dissatisfaction with existing peer review processes; to address bias in peer review; to subvert editorial selection processes that favour novelty over quality; to publish articles rejected by, or unsuitable for, journals. | *“Preprints have a huge psychological benefit for lab members as they see their work “published” while they go through the long, laborious process of peer-review.”*<br>*“It was too hard to fight with reviewers of journals”* | 12.6 | 10.8 |
| Feedback | To receive more feedback from peers before work is published in a journal; to allow studies to incorporate feedback and be updated with multiple versions before the final version is published. | *“Preprints can have versions, so you can publish version 1, without plans to submit to a journal yet (waiting on feedback from peers or coauthors).”*<br>*“For manuscripts reporting controversial results, I expected that depositing as preprints would allow gathering pre-submission feedback to better polish the finally submitted draft.”* | 7.1 | 8.1 |
| Open science | To provide free, unrestricted access to research results; to promote good open science practices and transparency; to support a public good; to reduce costs of journal publication. | *“I mostly just wanted to be able to provide my science to the public.”*<br>*“I believe its a principal duty for scientists to make their research available as open and free as possible.”* | 26.5 | 29.7 |
| Policies | To comply with policies and requests of journals, publishers, institutions (including individual departments) or funders requiring or suggesting that articles be deposited open access or on a preprint server. | *“It was a journal requirement.”*<br>*“While my institution does not require preprints, they want self-archiving, and preprints nicely do that.”* | 6.3 | 5.4 |
| Speed | To increase the speed until which research is available online; to reduce long delays in journal publication processes. | *“Speed of result dissemination - traditional publishing can be very slow (especially if more than one journal is tried in turn).”*<br>*“The knowledge that the submission, editor's review and peer-review process can take up to years.”* | 17.8 | 13.5 |
| Other | Labels that did not fit into one of the above categories, and was mentioned <10 times across all free-text responses for this question. | *“To have foundational information on our study sites citable for other publications based on research at those sites.”*<br>*“One of the preprints corresponds to a Spanish written article, so the visibility of an English vs Spanish paper changes, preprints allowed me to do so.”* | 7.1 | 16.2 |

Results from the free-text responses (**Table 4**) showed that career development is also an important motivation for submitting preprints (e.g. *“I was on the job market, so posting preprints was a way to show employers that I was productive and had products in publishing stages even though they weren’t yet in print.”*) for articles that participants served as corresponding authors (36.8% of all free-text responses mentioned this theme) and co-authors (29.7%). This includes submitting preprints to show evidence in job or grant applications, or simply to add research outputs to a CV that do not intend to be submitted to a journal. Other factors broadly mirrored those from the quantitative survey results: the next most-discussed theme was surrounding open science, i.e. posting preprints to support a free, unrestricted scientific system (e.g. *“Just seems the obviously correct thing to do, given that I basically want as many people to have (free) access to and read my work as possible.”;* 26.5% for corresponding authorships; 29.7% for co-authorships) and increasing the speed at which research is available (e.g. *“Move science forward quickly. We want to get our methods/findings/ideas out as quickly as possible so people can build off of them.”;* 17.8% for corresponding authorships; 13.5% for co-authorships). Many participants also discussed the themes relating to editorial processes, primarily expressing dissatisfaction with the current journal publishing system, including biases and a tendency to favour novelty over quality in peer review (e.g. *“Our, subjective, assessment was that our manuscript was being intentionally delayed/blocked in the peer-review process. Wanting to reach out with an important and timely message made us investigate the option of posting a preprint. “;* 12.6% for corresponding authorships; 10.8% for co-authorships). The potential to increase citation/online impact of work via preprints was also a well-discussed theme (e.g. *“I think that submitting some manuscripts as a preprint is a good idea because this will increase the possibility of citations before publishing in the scientific journals.”)*, although this was more common for articles that authors served as a corresponding author (20.2%) than as a co-author (5.4%). Other themes that motivated participants to post preprints were related to potential competition, receiving feedback, and policies, although all of these themes were discussed by less than 10% of participants for both corresponding authorships and co-authorships.

### Why do some authors not post preprints?

The previous section investigated factors that motivate authors to post their articles as preprints. In this section we explore the opposite view - factors that cause authors to not want to post their articles as preprints. In this section, survey participants who responded that they have posted either none or only some of their previously published journal articles as preprints (at the level of a corresponding author, and as a co-author) were presented with questions that focused specifically on the set of articles that were not posted as preprints. Results for the survey questions are shown in **Figure 7**, and regression results in **Table 5**; as with the previous section, participants were also provided with a free-text area to expand on these answers or add further reasoning (**Table 6**).

**Figure 7:**
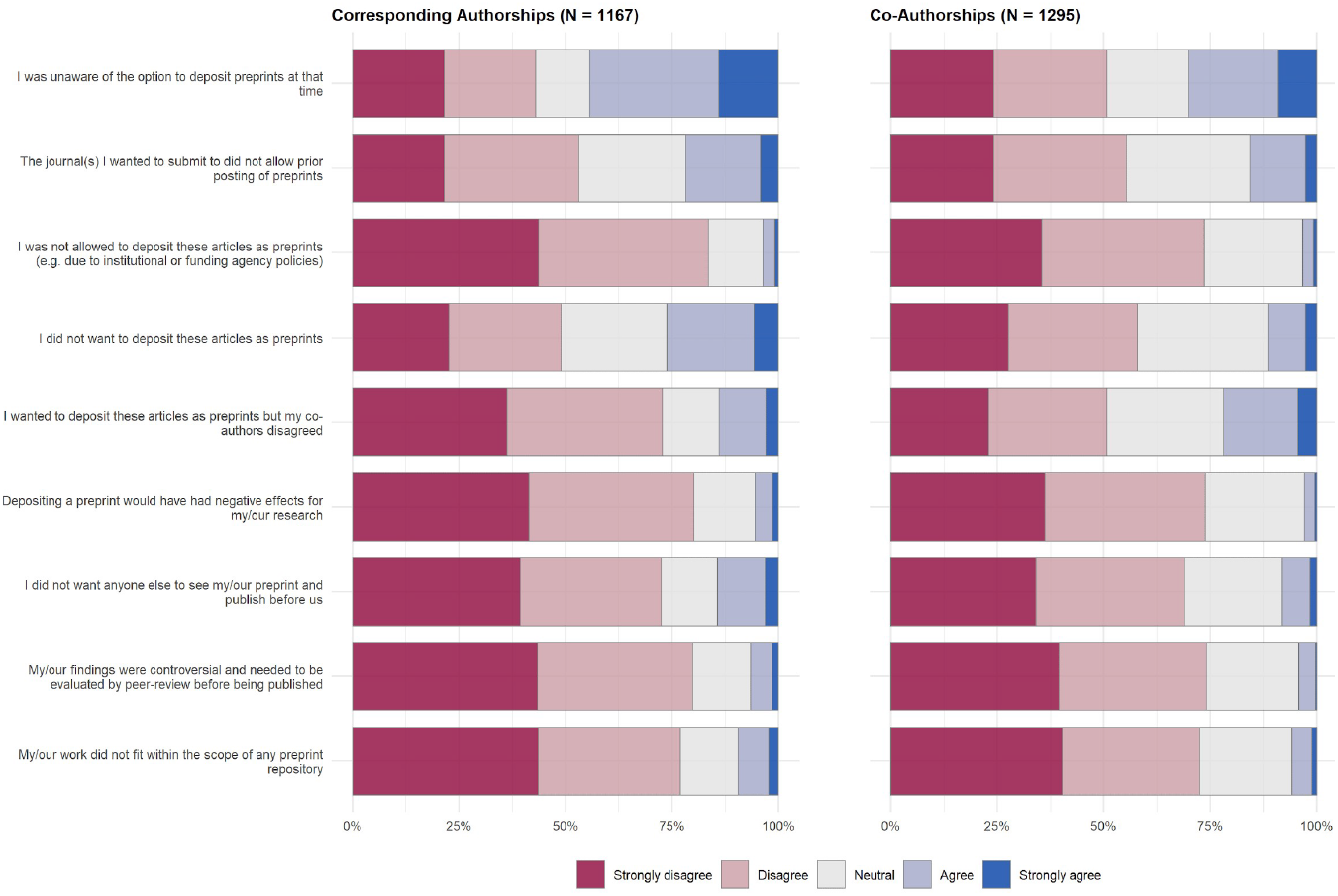
Reasons that survey participants did not post articles as preprints. Results show answers to survey questions on a 5-point Likert scale, divided by articles that authors published as a corresponding author (left) and articles that authors published as a co-author (right).

**Table 5:**
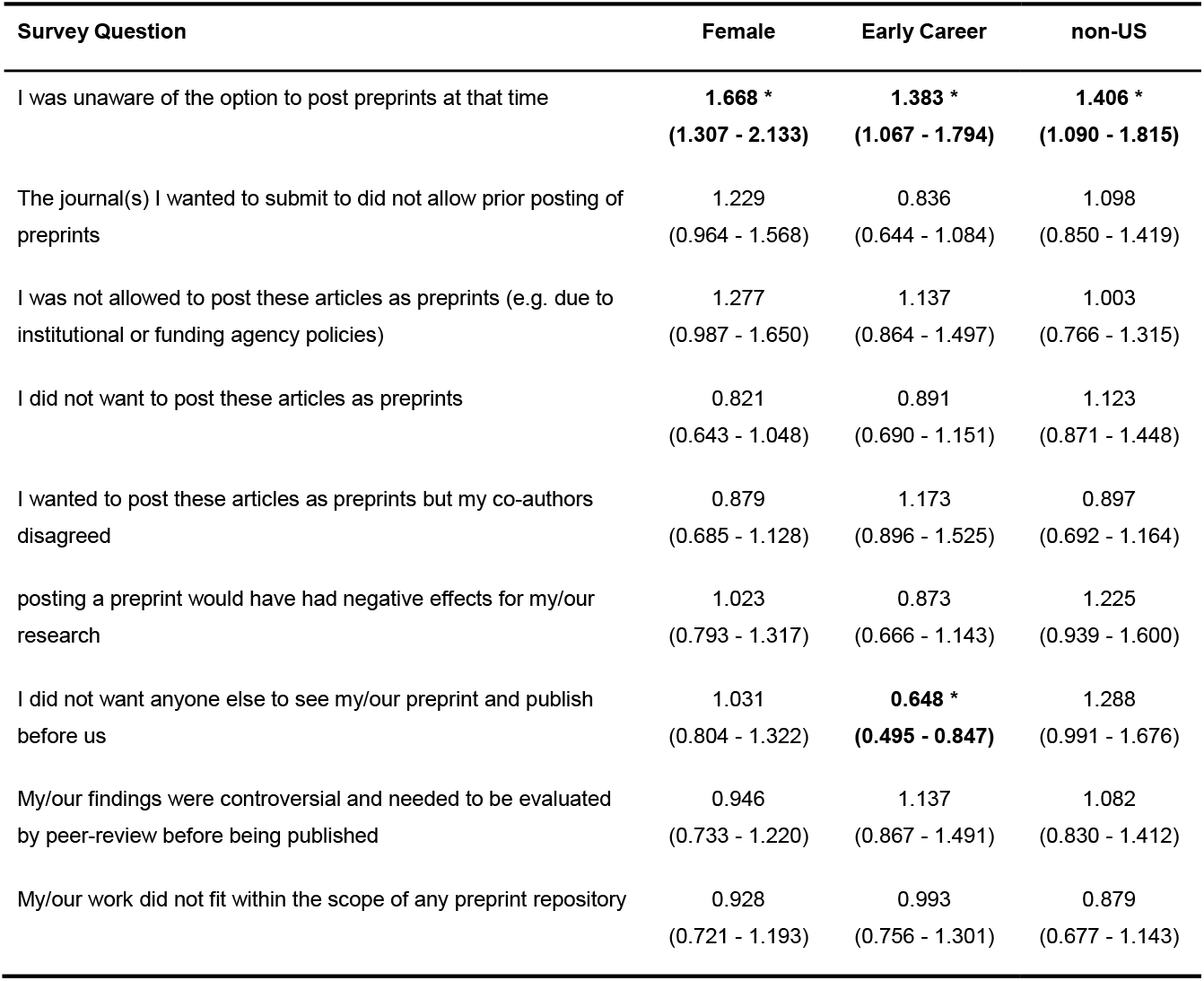
Results of ordinal logistic regression for survey questions in Figure 7 (corresponding authorships only; N = 1021). Values are presented as exponentiated odds-ratios, with the range in parentheses representing 95% confidence intervals. Significant results at the 95% level are indicated in **bold** and with “*”.

Overall, we found the strongest reason for authors to not post work as preprints to be that they were simply unaware of the option at the time that the work was conducted (44.4% agreed/strongly agreed for corresponding authorships, 30.1% for co-authorships). For the remainder of the questions, participants broadly disagreed with our potential reasons for not posting articles as preprints - in particular, participants disagreed/strongly disagreed that they were not allowed to post their articles as preprints (83.5% for corresponding authorships, 73.5% for co-authorships) or that posting a preprint would have had a negative effect on their work (80.2% for corresponding authorships, 73.8% for coauthorships). Interestingly, despite previous studies suggesting that the lack of quality assurance and premature reporting of incorrect/controversial results by the media are challenges for the establishment of preprints (ASAPbio, 2020; Chiarelli et al., 2019), in our results the majority of authors disagreed/strongly disagreed that their results were controversial and needed to be evaluated by peerreview before being published (79.8% for corresponding authorships, 74.2% for co-authorships).

Regression analysis for the set of questions presented in **Figure 7** are shown in **Table 5**. Female authors were more likely to report that they were unaware of the option to post preprints at the time their articles were published (OR: 1.668; 95% CI: 1.307 - 2.133) - the same was also true of early career researchers (OR: 1.383; 95% CI: 1.067 - 1.794) and non-US based researchers (OR: 1.406; 95% CI: 1.090 - 1.815). Early career researchers were also less likely to report that they did not post preprints because they did not want anyone else to see their preprint and publish before them (OR: 0.648; 95% CI: 0.495 - 0.847) - these results agree well with the previous section on motivations for posting preprints, where early career researchers were less likely to report that they post preprints to stake a priority claim on their findings, indicating overall that early career researchers are less motivated in their preprint publishing activities by competition compared to more senior scientists.

A summary of free-text responses elaborating on this section of the survey are shown in **Table 6**. Free-text responses confirmed some aspects of the qualitative survey results described above: participants reported most strongly that the reason for not posting articles that they served as corresponding author for was simply due to historic reasons, i.e. due to a lack of awareness of preprint servers or lack of existence of preprint servers at the time their journal articles were published (e.g. *“Before 2018, I was not familiar with preprint servers, and didn’t feel confident posting preprints simply because I wasn’t sure what to expect”;* 26.3% of corresponding authorships). Other important themes discussed regarding corresponding authorships were reluctance to post preprints prior to receiving feedback and quality control through journal peer review processes (e.g. *“To prevent people reading the paper before it had gone through rigorous peer review. One example would be the case of a paper with clinical findings. I wanted the manuscript to go through rigorous review before making it public in case we had made an error or unintentionally misrepresented some aspect of our findings.”*; 22.3% of corresponding authorships), article types that were not allowed or not suitable to be posted as preprints (e.g. *“In the field of taxonomy, I have the feeling that preprints may cause nomenclatural confusion. Therefore I would not opt for preprint deposition for alpha-taxonomic work, which consists part of my research.”;* 17.3% of corresponding authorships), and the extra labour and time required to format and submit preprints in addition to the journal formatting and submission process (e.g. *“Didn’t have time. Although it’s pretty easy, long author lists still take time to enter on bioRxiv.”;* 16.8% of corresponding authorships). Other lesser-discussed themes relating to corresponding authorships centred on journal policies that do not allow posting of preprints, disagreement from co-authors who did not want to post preprints, the lack of accessibility benefits when the article will be published in an open-access journal anyway, and the potential for preprints to lead to “scooping” of results by competing groups; although the latter factor was only discussed in a minority of comments (6.1% of corresponding authorships), it is interesting to note the conflict between authors who report that they post preprints as a method to prevent scooping and give them a competitive advantage, as noted in the previous section on preprint motivations, and authors who report that they do *not* post preprints to prevent scooping and retain their competitive advantage.

**Table 6:**
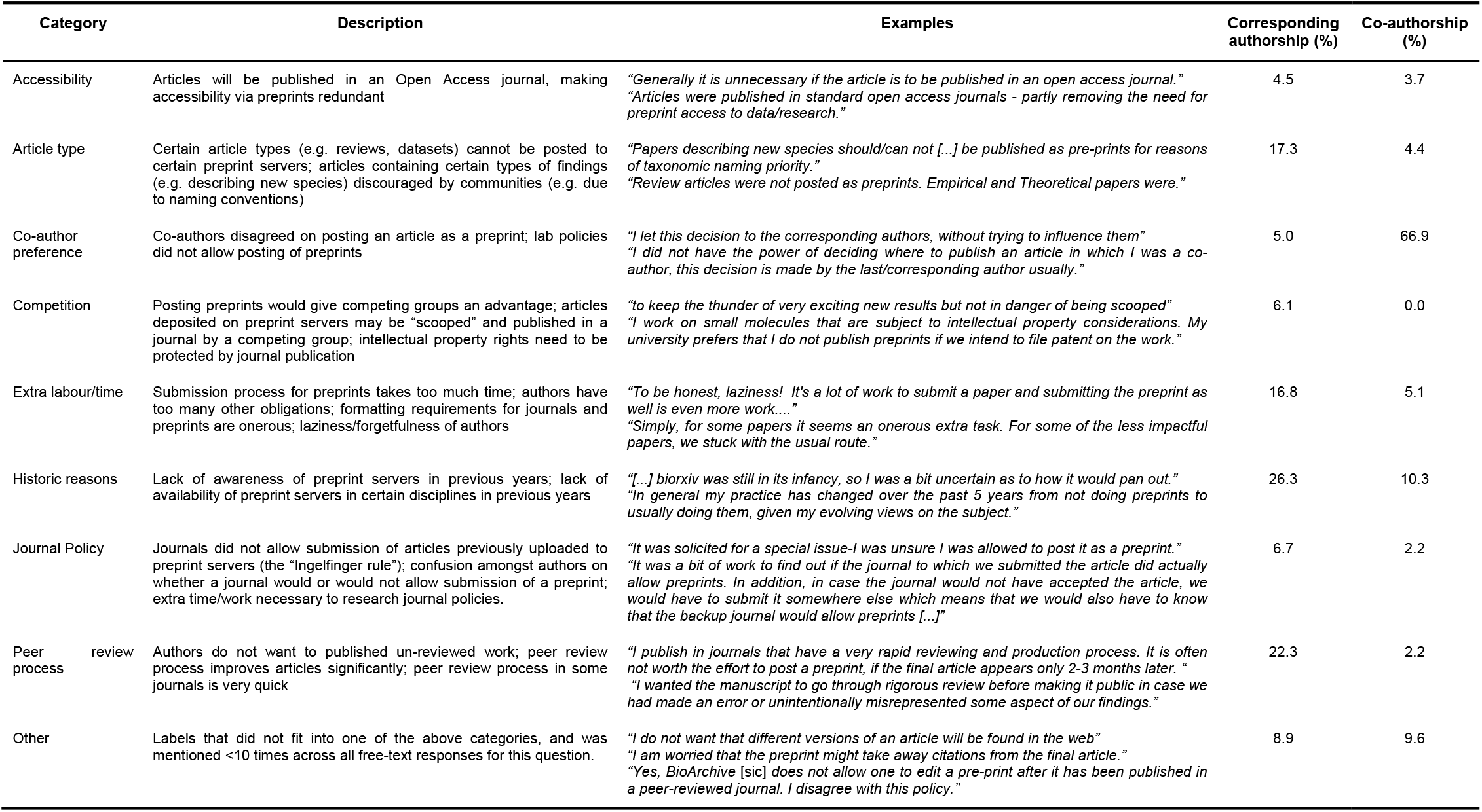
Classification of free-text responses, in response to the question “Were there any additional reasons that caused you to not deposit these articles as preprints?” Categories are ordered alphabetically (except for “Other”). Percentages for corresponding and co-authorships refer to the proportion of all responses containing the relevant category; responses could contain multiple categories (N = 180 comments on corresponding authorships, 137 comments on co-authorships).

With respect to co-authorships, by far the most discussed reason for not posting articles as preprints related to co-author preference (66.9% of free-text comments), underlining the importance of the corresponding author in deciding the publication strategy of an article.

### Differences between posted and non-posted preprints

Survey participants who only reported that they posted some of their previously published journal articles as preprints (**Figure 3**) were also asked to report on the differences between articles that they posted as preprints, and those that they did not. Results for the survey questions are shown in **Figure 8**, regression results in **Table 7** and additional free-text responses in **Table 8**.

**Figure 8:**
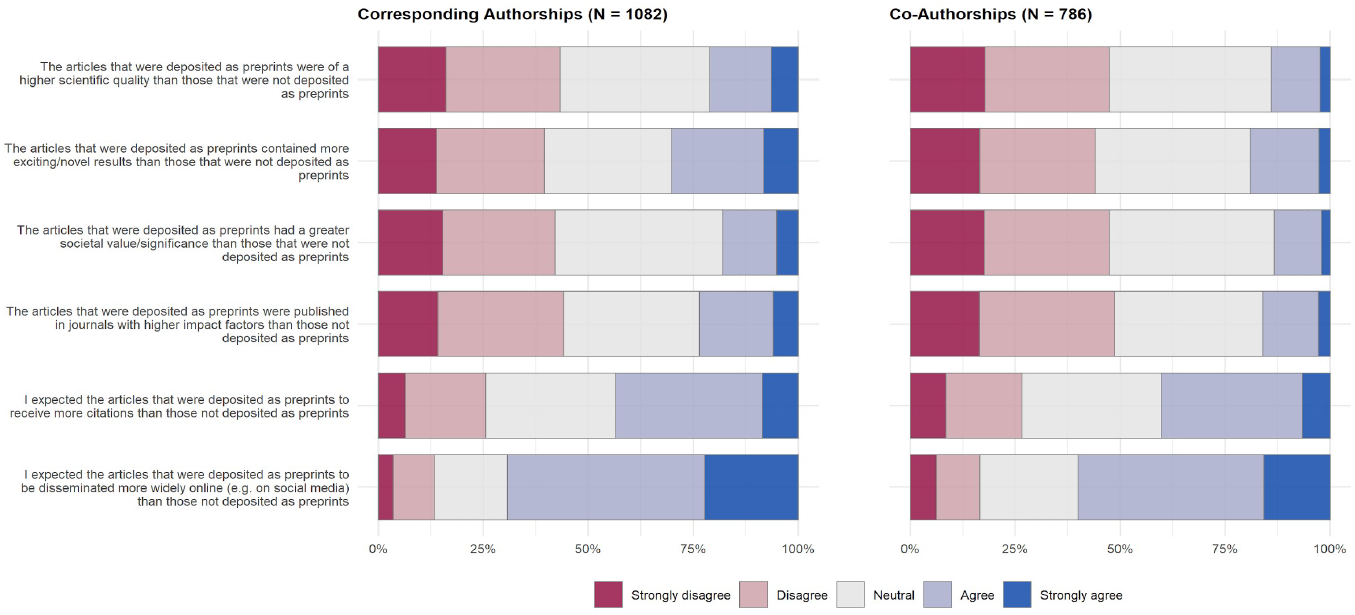
Differences between articles posted as preprints versus those not posted as preprints. Results show answers to survey questions on a 5-point Likert scale, divided by articles that authors published as a corresponding author (left) and articles that authors published as a co-author (right).

**Table 7:** Results of ordinal logistic regression for survey questions in Figure 8 (corresponding authorships only; N = 953). Values are presented as exponentiated odds-ratios, with the range in parentheses representing 95% confidence intervals. Significant results at the 95% level are indicated in **bold** and with “*”.

| Survey Question | Female | Early Career | non-US |
| --- | --- | --- | --- |
| The articles that were posted as preprints were of a higher scientific quality than those that were not posted | 0.826<br>(0.640 - 1.066) | 0.812<br>(0.618 - 1.067) | 1.049<br>(0.805 - 1.366) |
| The articles that were posted as preprints contained more exciting/novel results than those that were not posted | 0.834<br>(0.647 - 1.075) | 0.765<br>(0.584 - 1.002) | 1.026<br>(0.789 - 1.335) |
| The articles that were posted as preprints had a greater societal value/significance than those that were not posted | **0.687 \***<br>**(0.530 - 0.889)** | 0.906<br>(0.689 - 1.193) | 0.803<br>(0.612 - 1.053) |
| The articles that were posted as preprints were published in journals with higher impact factors than those not posted | 0.883<br>(0.683 - 1.141) | 0.816<br>(0.622 - 1.071) | 0.972<br>(0.746 - 1.266) |
| I expected the articles that were posted as preprints to receive more citations than those not posted | 0.836<br>(0.643 - 1.086) | 1.046<br>(0.797 - 1.374) | 1.097<br>(0.840 - 1.432) |
| I expected the articles that were posted as preprints to be disseminated more widely online than those not posted | 0.804<br>(0.614 - 1.052) | 1.082<br>(0.818 - 1.431) | 1.075<br>(0.816 - 1.413) |

**Table 8:**
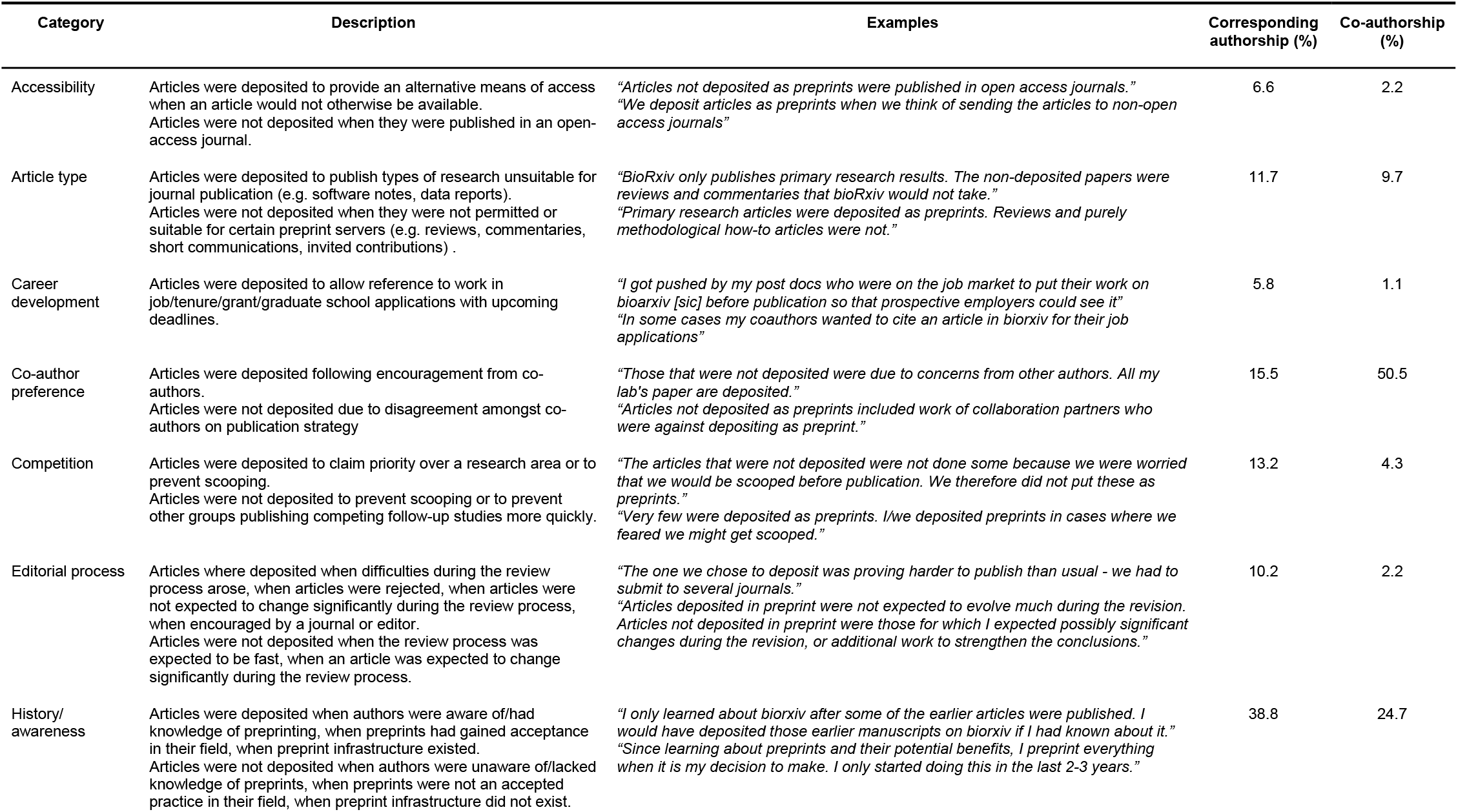
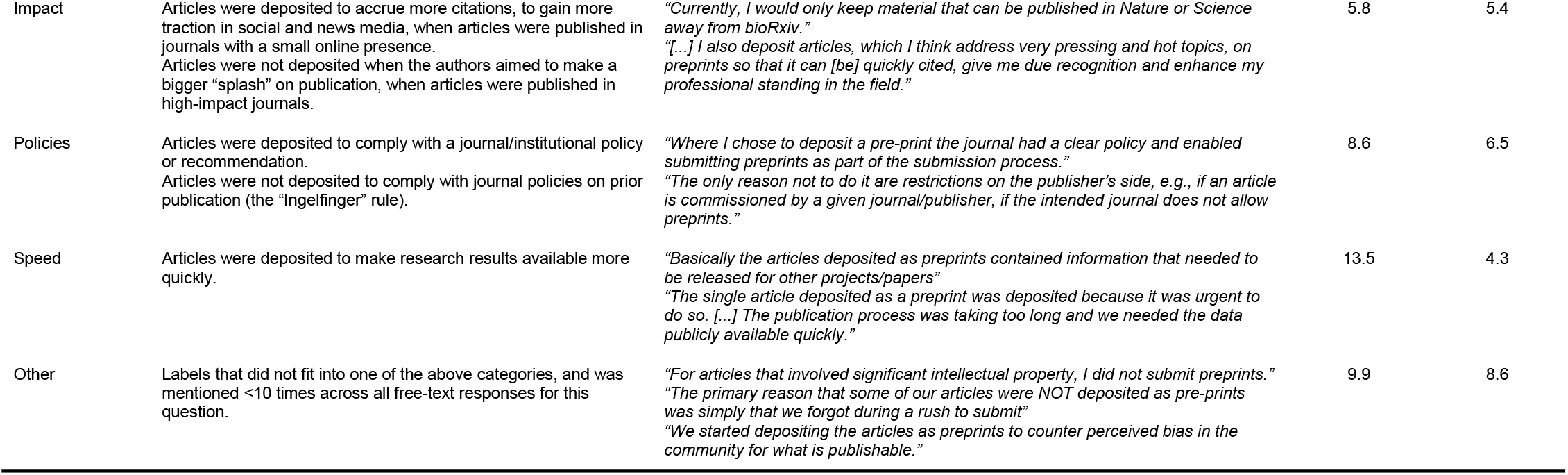
Classification of free-text responses, in response to the question “Were there any other differences between the articles you chose to deposit as preprints versus those you did not deposit, or differences in your expectations for how the articles would be received in your community?” Categories are ordered alphabetically (except for “Other”). Percentages for corresponding and co-authorships refer to the proportion of all responses containing the relevant category; responses could contain multiple categories (N = 394 comments on corresponding authorships, 93 comments on coauthorships).

| Category | Description | Examples | Corresponding authorship (%) | Co-authorship (%) |
| --- | --- | --- | --- | --- |
| Accessibility | Articles were deposited to provide an alternative means of access when an article would not otherwise be available.<br>Articles were not deposited when they were published in an open-access journal. | *“Articles not deposited as preprints were published in open access journals.”*<br>*“We deposit articles as preprints when we think of sending the articles to non-open access journals”* | 6.6 | 2.2 |
| Article type | Articles were deposited to publish types of research unsuitable for journal publication (e.g. software notes, data reports).<br>Articles were not deposited when they were not permitted or suitable for certain preprint servers (e.g. reviews, commentaries, short communications, invited contributions) . | *“BioRxiv only publishes primary research results. The non-deposited papers were reviews and commentaries that bioRxiv would not take.”*<br>*“Primary research articles were deposited as preprints. Reviews and purely methodological how-to articles were not.”* | 11.7 | 9.7 |
| Career development | Articles were deposited to allow reference to work in job/tenure/grant/graduate school applications with upcoming deadlines. | *“I got pushed by my post docs who were on the job market to put their work on bioarxiv [sic] before publication so that prospective employers could see it”*<br>*“In some cases my coauthors wanted to cite an article in biorxiv for their job applications”* | 5.8 | 1.1 |
| Co-author preference | Articles were deposited following encouragement from co-authors.<br>Articles were not deposited due to disagreement amongst co-authors on publication strategy | *“Those that were not deposited were due to concerns from other authors. All my lab’s paper are deposited.”*<br>*“Articles not deposited as preprints included work of collaboration partners who were against depositing as preprint.”* | 15.5 | 50.5 |
| Competition | Articles were deposited to claim priority over a research area or to prevent scooping.<br>Articles were not deposited to prevent scooping or to prevent other groups publishing competing follow-up studies more quickly. | *“The articles that were not deposited were not done some because we were worried that we would be scooped before publication. We therefore did not put these as preprints.”*<br>*“Very few were deposited as preprints. I/we deposited preprints in cases where we feared we might get scooped.”* | 13.2 | 4.3 |
| Editorial process | Articles where deposited when difficulties during the review process arose, when articles were rejected, when articles were not expected to change significantly during the review process, when encouraged by a journal or editor.<br>Articles were not deposited when the review process was expected to be fast, when an article was expected to change significantly during the review process. | *“The one we chose to deposit was proving harder to publish than usual - we had to submit to several journals.”*<br>*“Articles deposited in preprint were not expected to evolve much during the revision. Articles not deposited in preprint were those for which I expected possibly significant changes during the revision, or additional work to strengthen the conclusions.”* | 10.2 | 2.2 |
| History/ awareness | Articles were deposited when authors were aware of/had knowledge of preprinting, when preprints had gained acceptance in their field, when preprint infrastructure existed.<br>Articles were not deposited when authors were unaware of/lacked knowledge of preprints, when preprints were not an accepted practice in their field, when preprint infrastructure did not exist. | *“I only learned about biorxiv after some of the earlier articles were published. I would have deposited those earlier manuscripts on biorxiv if I had known about it.”*<br>*“Since learning about preprints and their potential benefits, I preprint everything when it is my decision to make. I only started doing this in the last 2-3 years.”* | 38.8 | 24.7 |
| Impact | Articles were deposited to accrue more citations, to gain more traction in social and news media, when articles were published in journals with a small online presence.<br>Articles were not deposited when the authors aimed to make a bigger “splash” on publication, when articles were published in high-impact journals. | *“Currently, I would only keep material that can be published in Nature or Science away from bioRxiv.”*<br>*“[...] I also deposit articles, which I think address very pressing and hot topics, on preprints so that it can [be] quickly cited, give me due recognition and enhance my professional standing in the field.”* | 5.8 | 5.4 |
| Policies | Articles were deposited to comply with a journal/institutional policy or recommendation.<br>Articles were not deposited to comply with journal policies on prior publication (the “Ingelfinger” rule). | *“Where I chose to deposit a pre-print the journal had a clear policy and enabled submitting preprints as part of the submission process.”*<br>*“The only reason not to do it are restrictions on the publisher’s side, e.g., if an article is commissioned by a given journal/publisher, if the intended journal does not allow preprints.”* | 8.6 | 6.5 |
| Speed | Articles were deposited to make research results available more quickly. | *“Basically the articles deposited as preprints contained information that needed to be released for other projects/papers”*<br>*“The single article deposited as a preprint was deposited because it was urgent to do so. [...] The publication process was taking too long and we needed the data publicly available quickly.”* | 13.5 | 4.3 |
| Other | Labels that did not fit into one of the above categories, and was mentioned <10 times across all free-text responses for this question. | *“For articles that involved significant intellectual property, I did not submit preprints.”*<br>*“The primary reason that some of our articles were NOT deposited as pre-prints was simply that we forgot during a rush to submit”*<br>*“We started depositing the articles as preprints to counter perceived bias in the community for what is publishable.”* | 9.9 | 8.6 |

Participants were initially asked to report on differences between articles in terms of article quality, novelty, societal value/significance, the impact factor of the publishing journal, as well as on their expectations for how articles would be received in terms of citations and other forms of online dissemination. Overall, a larger number of participants disagreed that articles they posted as preprints were higher quality (43.3% disagreed/strongly disagreed for corresponding authorships, 47.3% for coauthorships), more exciting/novel (39.6% disagreed/strongly disagreed for corresponding authorships, 44.0% for co-authorships), or had higher societal value/significance (42.1% disagreed/strongly disagreed for corresponding authorships, 47.4% for co-authorships). Participants also mainly disagreed that the articles they submitted as preprints were published in journals with higher impact factors than those not posts (44.1% disagreed/strongly disagreed for corresponding authorships, 48.7% for co-authorships).

Despite these results showing that authors did not preferentially post their highest quality, most novel or significant articles as preprints, authors still expected their articles to perform well in terms of impact: 43.5% of corresponding authorships and 40.2% of co-authorships agreed/strongly agreed that they expected their articles they posted as preprints to receive more citations than the ones they did not posted, with the proportions rising to 69.3% and 60.1% for corresponding authorships and coauthorships, respectively, that expected articles they posted as preprints to be disseminated more widely online than those not posted. Ostensibly, these results are at odds with each other: on one hand authors report that there are no differences in quality/novelty/significance of articles that they choose to post as preprints, and yet they still expect that the articles they posted will perform better in terms of citations and online dissemination.

With respect to our regression analysis, overall we find little difference in reported results between different demographic groups, although female authors were less likely to report that they post articles as preprints that were expected to have a greater societal value/significance than those that were not posted (OR: 0.687, 95% CI: 0.530-0.889) (**Table 7**).

Free-text responses **(Table 8)** detailed a range of additional differences that authors report between articles they do and do not post as preprints. With respect to corresponding authorships, 38.8% of respondents pointed to history/awareness of preprints as a reason for only posting a proportion of their articles as preprints, i.e. articles that they did not post as preprints were simply due to not knowing about preprints, or lack of establishment of preprints as an accepted norm at the time that they published their non-posted articles (e.g. *“the main reason for not publishing on a preprint server* was *that this was not very common in the field (Biology/biophysics) a few years ago”)*, Other factors that drove corresponding authors to post certain papers were related to co-author preference (e.g. *“a major criterion was on whether all authors agreed on posting them as preprint. As soon as one author objected, they were not posted”;* 15.5% of corresponding authorships), speed (e.g. *“The single article posted as a preprint was posted because it was urgent to do so. The data were useful for Endangered Species Act listing as well as for policies on invasive species mitigation. The publication process was taking too long and we needed the data publicly available quickly”;* 13.5% of corresponding authorships), and competition (e.g. *“The articles posted as preprint[sic] were more at risk of been scooped”;* 13.2% of corresponding authorships). With respect to competition, we found conflicting reasoning for posting or not posting certain articles as preprints: some respondents considered preprints as a way to gain a competitive advantage by claiming priority over a research finding (“*Articles posted were in possible competition, so ensured to be out first*”), whilst others regarded the posting of preprints a competitive disadvantage (“*Other novel results were not posted […] to protect from competing groups to overtake with follow-ups before we could finish ours*”). In terms of coauthorships, by far the most important factor that determined whether certain preprints were posted was co-author preferences (e.g. *“For articles where I was not corresponding author I usually had a much more limited role, and I did not take part in deciding where to submit the articles (as preprints or for journals)”*; 50.5% of co-authorships): these results are in good agreement with our previous section on decision-making for preprint posting **(Figure 4)**, where the majority of corresponding authors reported that they made the decision themselves to post an article as a preprint.

## Discussion and Conclusions

We used a mixed-methods survey approach to investigate the motivations, concerns and selection biases of authors in posting preprints to bioRxiv, a preprint server for the biological sciences. These results are important to inform bibliometric studies aiming to better understand the causal effects of preprint publishing models on citations or other metrics of dissemination (i.e. altmetrics), as well as to better understand the diffusion of preprints into different communities, which may further inform future policies of scholarly stakeholders including researchers, institutions, funders and providers of preprint infrastructure.

At the basic level, our results confirm results from previous surveys and interviews that have investigated the factors that motivate and prevent authors from posting preprints (e.g. ASAPbio, 2020; Chiarelli et al., 2019; Foster et al., 2017; Kelly, 2018; Sever et al., 2019): we find the strongest motivations to post preprints are to increase awareness of research and share findings more quickly (**Figure 5**), although we find some differences between demographic groups, e.g. early career researchers appear to be more strongly driven to post preprints to increase awareness of their research and to receive feedback compared to late career researchers, who were more strongly motivated to post preprints to stake a priority claim on their work (**Table 2**). An interesting aspect of these results, is that whilst survey participants were generally highly motivated to post preprints to increase awareness of their research, agreement was less strong on whether posting preprints actually had a positive benefit in terms of citations and/or metrics of online sharing (**Figure 6**), e.g. only 38.7% of authors agreed/strongly agreed that posting preprints had a positive effects on citations to their work. Such findings are highly relevant for studies aiming to understand the “true” citation effect of posting preprints. Qualitative results derived from free-text comments (**Table 4**) supported the quantitative survey results, but added additional dimensions and reasons that were not previously captured, e.g. the high importance of preprints for career development and high levels of dissatisfaction with current journal publishing systems.

We additionally investigated the factors that cause authors to not publish articles as preprints, revealing a mixture of structural and self-determined reasons that preprints were not posted (**Figure 7; Tables 5, 6**). In particular, both our quantitative and qualitative results show that lack of awareness of preprints plays the most important role in their posting, with other important factors including hesitation to publish work that has not undergone peer review, the extra time and effort required to format and submit preprints, and the discouragement of publishing certain types of articles as preprints (e.g. bioRxiv do not allow publishing of review articles on their platform^2^). Although the so-called “Ingelfinger rule” is often mentioned as a factor that discourages authors from posting preprints, we found that journal policies only affected a minority of participants decisions to post preprints, in line with findings from a similar survey from ASAPbio (2020) that found that external pressures rank relatively low on researchers motivations and concerns. A small number of participants also reported that confusion in whether a journal would or would not accept a preprint discouraged them from posting a preprint, which may be rooted in the high proportion of journals that still do not have clear preprint policies (Klebel et al., 2020).

A novel aspect of our survey in comparison to previously-mentioned studies is that we directly asked authors who have only posted a subset of their published journal articles as preprints about the differences between those they did and did not post. From the quantitative survey results (**Figure 8**), we found that participants generally disagreed that articles chosen and not chosen to be posted as preprints differed in their quality, novelty or societal value/significance. Despite this, 43.5% of participants agreed/strongly agreed that they expected that their articles posted as preprints would receive more citations than those not posted as preprints, and 69.3% agreed/strongly agreed that they expected that their articles posted as preprints would be shared more widely online than those not posted as preprints. These results appear to be in conflict with the first dimension of the Self-Selection Bias postulate, as previously discussed (i.e. that authors preferentially select their highest quality/most novel/most significant to post as preprints), - the majority of authors do not appear to be actively selecting articles to post as preprints based on subjective criteria. A question therefore remains at why authors still expect their articles to receive more citations/be shared more widely online even if they do not believe those articles to differ in terms of quality/novelty/significance - the answer may lie in their motivations for submitting preprints in the first place, i.e. to increase awareness of their work and share findings more quickly, which could be expected to drive more citations and increase metrics of online sharing.

In summary, our results suggest that previous findings of a citation/altmetric advantage for bioRxiv preprints (e.g. Serghiou & Ioannidis, 2018; Fu & Hughey, 2019; Fraser et al., 2020) are not strongly influenced by selection biases of preprint authors. However, we do not rule out that biases exist amongst the authors of preprints themselves, e.g. preprints may be preferentially authored by those “highest quality” researchers, or researchers who tend to work on more novel subject areas where early dissemination of results has a stronger effect than other less novel areas. In the current study we have not investigated this additional dimension of the Self-Selection Bias postulate, and encourage future studies in this area.

### Limitations

We acknowledge a number of important limitations of our study, which may be improved and built upon in future studies. Most importantly, we use a survey approach which requires self-reporting from respondents on their preprinting activities. Thus, we must make the assumption that any responses are reported honestly and without bias. During the survey we asked authors about their own potential biases in selecting articles that they have previously posted as preprints, versus those that were not posted as preprints. However, in distributing our survey we may also introduce bias amongst survey participants - firstly that we have targeted primarily corresponding authors of preprints on bioRxiv (as we collected email address of bioRxiv preprint authors), and secondly that those who responded to the survey are potentially authors who are more engaged with preprints in general. Future studies may therefore consider targeting authors of journal articles who have never posted preprints, to determine if differences exist for authors who are less engaged with, or knowledge about, preprints. Whilst we have made efforts to understand the demographics of our survey respondents and the influence of these different groups on our survey results, there exist some biases in our sample (e.g. the overrepresentation of US-based and male authors) that mean that we should be cautious about generalising findings to the wider scientific system.

Our survey was conducted in March and April 2020, and targeted authors of preprints that were posted between November 2013 and December 2018. The use and visibility of preprints has been growing in recent years, and the last year in particular has seen a surge of preprints published in response to the COVID-19 pandemic (Fraser et al., 2021). Long-term monitoring and/or future replication of our study will be necessary to understand how authors’ preprinting behaviour evolves over time, and what the long-term effects of “shocks” to the scientific publishing system, such as COVID-19, will have on the future development and usage of preprints.

Lastly, we focus our study on a single preprint server for biological sciences. We encourage additional studies that focus on, and compare our results with, other research disciplines where the available preprint infrastructure (e.g. discipline-specific preprint servers) and usage amongst researchers differs.

## Acknowledgements

This work is supported by BMBF project OASE, grant numbers 01PU17005A and 01PU17005B. We thank Athanasios Mazarakis for support in managing the LimeSurvey server through which this survey was conducted.

## Author Contributions

Conceptualisation: NF, PM, IP

Data curation: NF

Formal analysis: NF

Funding acquisition: PM, IP

Investigation: NF

Methodology: NF

Project administration: PM, IP

Resources: PM, IP

Software: NF

Supervision: PM, IP

Validation: NF

Visualisation: NF

Writing - original draft: NF

Writing - review and editing: NF, PM, IP

## Data and Code Availability

Survey templates, response data and all code used for the preparation, analysis and visualisation of response data are available on GitHub (https://github.com/nicholasmfraser/biorxiv_survey/) and archived on Zenodo (https://zenodo.org/10.5281/zenodo.5166749). Note that raw free-text responses and email addresses of survey respondents were removed to preserve participant anonymity.

## Footnotes

1 The “Ingelfinger rule” stems from an editorial piece in the New England Journal of Medicine, written by then-Editor Franz Ingelfinger (‘Definition of Sole Contribution’, 1969). The editorial documented a case of a manuscript submitted to the journal that had previously appeared in print elsewhere; the journal rejected the manuscript and implemented a policy that manuscripts submitted to the journal must not be published nor submitted elsewhere.

2 See https://www.biorxiv.org/about/FAQ

## References

Abdill, R. J., & Blekhman, R. (2019). Tracking the popularity and outcomes of all bioRxiv preprints. ELife, 8, e45133. https://doi.org/10.7554/eLife.45133

Archambault, É., Côté, G., Struck, B., & Voorons, M. (2016). Research impact of paywalled versus open access papers. Copyright, Fair Use, Scholarly Communication, Etc. https://digitalcommons.unl.edu/scholcom/29/

ASAPbio. (2020). Preprint authors optimistic about benefits: Preliminary results from the #bioPreprints2020 survey. https://asapbio.org/biopreprints2020-survey-initial-results

Björk, B.-C., & Solomon, D. (2013). The publishing delay in scholarly peer-reviewed journals. Journal of Informetrics, 7(4), 914–923. https://doi.org/10.1016/j.joi.2013.09.001

Chamberlain, S., Zhu, H., Jahn, N., Boettiger, C., & Ram, K. (2020). rcrossref: Client for Various ‘CrossRef’ ‘APIs’. https://CRAN.R-project.org/package=rcrossref

Chiarelli, A., Johnson, R., Pinfield, S., & Richens, E. (2019). Preprints and Scholarly Communication: An Exploratory Qualitative Study of Adoption, Practices, Drivers and Barriers. F1000Research, 8, 971. https://doi.org/10.12688/f1000research.19619.2

Cobb, M. (2017). The prehistory of biology preprints: A forgotten experiment from the 1960s. PLOS Biology, 15(11), e2003995. https://doi.org/10.1371/journal.pbio.2003995

Davis, P. M., & Fromerth, M. J. (2007). Does the arXiv lead to higher citations and reduced publisher downloads for mathematics articles? Scientometrics, 71(2), 203–215. https://doi.org/10.1007/s11192-007-1661-8

Definition of Sole Contribution. (1969). New England Journal of Medicine, 281(12), 676–677. https://doi.org/10.1056/NEJM196909182811208

Delamothe, T., Smith, R., Keller, M. A., Sack, J., & Witscher, B. (1999). Netprints: The next phase in the evolution of biomedical publishing. BMJ, 319(7224), 1515–1516. https://doi.org/10.1136/bmj.319.7224.1515

Foster, J., Hearst, M., Joakim, N., & Shiqi, Z. (2017). Report on ACL Survey on Preprint Publishing and Reviewing. https://www.aclweb.org/portal/sites/default/files/SurveyReport2017.pdf

Fraser, N., Brierley, L., Dey, G., Polka, J. K., Pálfy, M., Nanni, F., & Coates, J. A. (2021). The evolving role of preprints in the dissemination of COVID-19 research and their impact on the science communication landscape. PLOS Biology, 19(4), e3000959. https://doi.org/10.1371/journal.pbio.3000959

Fraser, N., Momeni, F., Mayr, P., & Peters, I. (2020). The relationship between bioRxiv preprints, citations and altmetrics. Quantitative Science Studies, 1–21. https://doi.org/10.1162/qss_a_00043

Fu, D. Y., & Hughey, J. J. (2019). Releasing a preprint is associated with more attention and citations for the peer-reviewed article. ELife, 8, e52646. https://doi.org/10.7554/eLife.52646

Gargouri, Y., Hajjem, C., Larivière, V., Gingras, Y., Carr, L., Brody, T., & Harnad, S. (2010). Self-Selected or Mandated, Open Access Increases Citation Impact for Higher Quality Research. PLoS ONE, 5(10), e13636. https://doi.org/10.1371/journal.pone.0013636

Ginsparg, P. (2011). ArXiv at 20. Nature, 476(7359), 145–147. https://doi.org/10.1038/476145a

Johansson, M. A., Reich, N. G., Meyers, L. A., & Lipsitch, M. (2018). Preprints: An underutilized mechanism to accelerate outbreak science. PLOS Medicine, 15(4), e1002549. https://doi.org/10.1371/journal.pmed.1002549

Kelly, D. (2018). SIGIR Community Survey on Preprint Services. ACM SIGIR Forum, 52(1), 11–33. https://doi.org/10.1145/3274784.3274787

Kirkham, J. J., Penfold, N., Murphy, F., Boutron, I., Ioannidis, J. P., Polka, J. K., & Moher, D. (2020). A systematic examination of preprint platforms for use in the medical and biomedical sciences setting. BioRxiv. https://doi.org/10.1101/2020.04.27.063578

Klebel, T., Reichmann, S., Polka, J., McDowell, G., Penfold, N., Hindle, S., & Ross-Hellauer, T. (2020). Peer review and preprint policies are unclear at most major journals. PLOS ONE, 15(10), e0239518. https://doi.org/10.1371/journal.pone.0239518

Larivière, V., Sugimoto, C. R., Macaluso, B., Milojević, S., Cronin, B., & Thelwall, M. (2014). arXiv Eprints and the journal of record: An analysis of roles and relationships: arXiv E-Prints and the Journal of Record. Journal of the Association for Information Science and Technology, 65(6), 1157–1169. https://doi.org/10.1002/asi.23044

Moed, H. F. (2007). The effect of “open access” on citation impact: An analysis of ArXiv’s condensed matter section. Journal of the American Society for Information Science and Technology, 58(13), 2047–2054. https://doi.org/10.1002/asi.20663

National Institutes of Health. (2017). Reporting Preprints and Other Interim Research Products (Notice Number: NOT-OD-17-050). https://grants.nih.gov/grants/guide/notice-files/not-od-17-050.html

Nature. (2007). Community service. Nature, 447(7145), 614–614. https://doi.org/10.1038/447614a

Nature. (2012). https://www.nature.com/content/npg/23909.html

Penfold, N. C., & Polka, J. K. (2020). Technical and social issues influencing the adoption of preprints in the life sciences. PLOS Genetics, 16(4), e1008565. https://doi.org/10.1371/journal.pgen.1008565

Piwowar, H., Priem, J., Larivière, V., Alperin, J. P., Matthias, L., Norlander, B., Farley, A., West, J., & Haustein, S. (2018). The state of OA: A large-scale analysis of the prevalence and impact of Open Access articles. PeerJ, 6, e4375. https://doi.org/10.7717/peerj.4375

R Core Team. (2020). R: A Language and Environment for Statistical Computing. R Foundation for Statistical Computing. https://www.R-project.org/

Serghiou, S., & Ioannidis, J. P. A. (2018). Altmetric Scores, Citations, and Publication of Studies Posted as Preprints. JAMA, 319(4), 402. https://doi.org/10.1001/jama.2017.21168

Sever, R., Roeder, T., Hindle, S., Sussman, L., Black, K.-J., Argentine, J., Manos, W., & Inglis, J. R. (2019). bioRxiv: The preprint server for biology. BioRxiv. https://doi.org/10.1101/833400

Tennant, J., Bauin, S., James, S., & Kant, J. (2018). The evolving preprint landscape: Introductory report for the Knowledge Exchange working group on preprints. (MetaArXiv) [Preprint]. MetaArXiv. https://doi.org/10.31222/osf.io/796tu

Thomas, D. (2006). A General Inductive Approach for Analyzing Qualitative Evaluation Data. American Journal of Evaluation, 27(2), 237–246. https://doi.org/10.1177/1098214005283748

Varmus, H. (1999). E-Biomed: A Proposal for Electronic Publications in the Biomedical Sciences (Draft and Addendum). National Institutes of Health. https://profiles.nlm.nih.gov/spotlight/mv/catalog/nlm:nlmuid-101584926X356-doc

Venables, W. N., & Ripley, B. D. (2002). Modern Applied Statistics with S (Fourth). Springer. http://www.stats.ox.ac.uk/pub/MASS4

Wang, Z., Chen, Y., & Glänzel, W. (2020). Preprints as accelerator of scholarly communication: An empirical analysis in Mathematics. Journal of Informetrics, 14(4), 101097. https://doi.org/10.1016/j.joi.2020.101097

Wang, Z., Glänzel, W., & Chen, Y. (2020). The impact of preprints in Library and Information Science: An analysis of citations, usage and social attention indicators. Scientometrics, 125(2), 1403–1423. https://doi.org/10.1007/s11192-020-03612-4

Wickham, H. (2020). rvest: Easily Harvest (Scrape) Web Pages. https://CRAN.R-project.org/package=rvest

